# Human Satellite III long non-coding RNA imparts survival benefits to cancer cells

**DOI:** 10.1101/2021.09.30.462592

**Authors:** Manjima Chatterjee, Sonali Sengupta

## Abstract

Long non-coding RNAs are heterogeneous group of transcripts that lack coding potential and have crucial roles in gene regulations. Recent days have seen an increasing association of non-coding RNAs with human diseases, especially cancers. Satellite III (SatIII) lncRNAs are transcribed from pericentromeric heterochromatic region of the human chromosome. Though transcriptionally silent in normal conditions, SatIII is actively transcribed under condition of stress, mainly heat shock. SatII repeat, another component of pericentromeric region of human chromosome, has been associated with wide variety of epithelial cancer. Overexpression of Satellite RNAs induces breast cancer in mice. Though much is known about Satellite RNAs, which includes alpha satellites and SatII repeats, however little is known about SatIII in human cancers. Hence we directed our study to understand the role of human Satellite III repeats in cancerous conditions. In the present study, we show that colon and breast cancer cells transcribe SatIII independent of heat shock, in an HSF1-independent manner. Our study also reveals that, overexpression of SatIII RNA favours cancer cell survival by overriding chemo drug-induced cell death. Knockdown of SatIII sensitizes cells towards chemotherapeutic drugs. SatIII transcript knockdown restores the expression of p53 protein, which in turn facilitates cell death. Heat shock however helps SatIII to continue with its pro-cell survival function. Our results, therefore suggest SatIII to be an important regulator of human cancers. Induction of SatIII is not only a response to the oncogenic stress but also facilitates cancer progression by a distinct pathway that is different from heat stress pathway.

## 1. INTRODUCTION

Non-coding RNAs are transcripts that do not code for any protein. They are highly abundant in our genome and are important for cellular processes like chromatin remodelling, transcription, splicing, genome integrity etc. Non-coding RNAs are broadly classified into short and long non-coding RNAs based on their sizes. Short non-coding RNAs are small RNAs lesser than 200 bp long and includes siRNA and miRNA important for gene silencing. Long noncoding RNAs are RNAs more than 200 bp and they can extend up to several kilobases long (*reviewed in* Bartonicek et al., 2016; Lakhotia 2016). Long non-coding RNAs are also characterized according to their localization in human cells, structure, function, and interaction with DNA elements, protein-coding genes or other abundant RNAs. LncRNAs in general, function as molecular signals; they may scaffold protein complexes, or release proteins from chromatin as decoys (Bolha et al., 2017). Long non-coding RNAs play major roles in human diseases including neurodegenerative diseases and cancers (*reviewed in* Sengupta, 2017; Chatterjee & Sengupta, 2019). In cancer, they are tissue specific and are tightly regulated in association with genes responsible for cell cycle regulation, immune response and pluripotency (Bartonicek et al., 2016). Therefore, the lncRNAs are functionally important in defining fate of a pre-cancerous cell and may confer both oncogenic and tumor suppressing activity (Beckedorff et al., 2013). They are also well-known for guiding ribonucleoprotein (RNP)-complexes to specific gene targets as *cis* (neighbouring genes) or *trans* (distantly located genes) regulators (Lakhotia et al., 2020). The human SatIII lncRNA is a 2 to >5 kb transcript, that during heat shock, is transcribed by the heat shock factor 1 (HSF1) from the pericentromeric heterochromatic locus of chromosome 9q12 (Jolly & Lakhotia, 2006). Its transcription is directed by RNA polymerase II, polyadenylated, variable in size, however, contains a characteristic GGAAT motif of a typical satellite III repeat (Jolly et al., 2004). The HSF1-activated SatIII RNAs accumulate at the site of transcription and produce granule-like structures known as the nuclear stress bodies (nSBs). Besides its colocalization with HSF1, the SatIII-nSBs also recruit several splicing factors like, serine-arginine rich proteins ASF/SF2, SRp30c and 9G8, transcription factors like the CREB binding protein (CREBBP) and other hnRNPs (Jolly et al., 2004; Metz et al., 2004; Sengupta et al., 2009; Goenka et al., 2016). A recent study has identified 141 proteins as putative components of the SatIII associated nuclear stress bodies; many of these appeared to be nuclear RNA-binding proteins having functions in pre-mRNA splicing and processing (Ninomiya et al., 2020).

The heat exposed cells exhibit a highly conserved self-defense heat shock response (HSR) mechanism, which results into rapid induction of a group of chaperones, known as the heat shock proteins (HSPs). This provides an immediate protection against stress-induced damage (Lindquist & Craig, 1988). The role of human SatIII lncRNA in cellular recovery from stress is thus well narrated in earlier studies, but its specific involvement in cancer is yet to be unveiled. A few among several well-studied cancer specific lncRNAs are HOTAIR, MALAT-1, PANDA, PVT-1, lincRNA-p21, GAS5, MEG3 etc. They may act as oncogenes or tumor suppressors, and regulate a series of oncogenic events. Other than a key regulator of HSR, the HSF1 also possesses oncogenic potential and effectively permits cells to survive proteotoxic stress (Dai & Sampson, 2016). In cancer cells, the HSF1 remains constitutively active and maintains malignant phase as efficiently as it promotes tumor growth (Dai & Sampson, 2016). Condition like hyperosmotic stress induces SatIII independent of HSF1, rather, involves the tonicity enhancer-binding protein (TonEBP) as a transcription factor. This therefore suggests SatIII may involve different transcriptional factors for its expression under various growth conditions (Valgardsdottir et al., 2008).

Pericentromeric region of the human chromosome contains several satellite repeats, such as Human Satellite II and Human Satellite III. A very important feature of these satellite repetitive sequences is that, they are rarely transcribed under normal homeostatic conditions. Overexpression of satellite sequences has been associated with genomic instability. Ting et al. (2011) showed that there was a 40% overexpression of pericentromeric satellite repeats in mouse pancreatic ductal adenocarcinoma cells (PDACs) as compared to normal tissue. The same phenomenon was observed in human tissues as the satellite transcripts showed 21% overexpression in PDACs as compared to normal tissues. Further, Zhu et al. (2011) showed, satellite DNAs overexpress in a variety of human cancer including BRCA1 mutant cells. However, all this studies were limited to Human Satellite II repeats. It was important and intriguing at this point to investigate the role of Satellite III repeats in human cancer.

It is interesting to note that other than heat shock, a series of stress-enhancing chemicals also induce SatIII from multiple transcriptional loci as demonstrated in an early report by Sengupta et al. (2009). Chemicals that enhance HSP70 levels such as 8-hydroxyquinoline, zinc sulfate and ibuprofen are few of them that potentially induce SatIII similar to that of the heat induced-nSBs (Sengupta et al., 2009). A likely observation reveals treatment of cells with chemicals (MG132, lactacystin and puromycin) that enhance proteotoxic load are also capable of inducing SatIII-nSBs in a stress specific manner (Sengupta et al., 2009). As our study focuses on the expression of SatIII in context of cancer, we selected a series of anti-tumorigenic drugs that majorly target the stages of cell cycle. It is interesting to note that the anti-cancer drugs that function as effective stressors to kill tumor cells may themselves trigger adaptive stress response to the tumor environment to overcome on-going DNA damage repair (Tiligada, 2006). Staurosporine is an example of such a potent anti-neoplastic drug that induces apoptosis via G_2_/M phase arrest, and inhibits CDKs and protein kinase C available abundantly in the tumor niche (Antonsson & Persson, 2009). We then selected 5-fluorouracil (5-FU) that induces apoptosis in a caspase dependent pathway; it also targets the RNA bases and regulates growth metabolism to exert cytotoxic effects on the dividing cells (Mhaidat et al., 2014). Further we also included cisplatin (cis-Diamminedichloroplatinum (II): CDDP), known to introduce inter- and intra-strand crosslinked DNA-breaks to induce apoptosis in the breast cancer cells (Tanida et al., 2012). Notably, these drugs are able to moderate expressions of many lncRNAs (Teschendorff et al., 2015), while some of the lncRNA transcripts induce chemo-resistance to the cancer cells (*reviewed in* Zhang et al., 2020). Importance of p53 on cellular sensitivity to anti-cancer agents has been a topic of interest since decades. In some cases, inactivation of p53 leads to increased resistance indicating pathways leading to cell death are p53 dependent. While the other cases involve disruption of wild-type p53 function leading to increased drug sensitivity (*reviewed in* Ferreira et al., 1999). Therefore the protein has considerable role in upregulating genes involved in cell cycle regulation and apoptosis (Fischer, 2017). Interestingly, many long non-coding RNAs are part of the transcriptome and regulate chemo-resistance through p53 apoptotic signaling pathways (Deng et al., 2017; Hu et al., 2018).

Crucial role of SatIII in HSR is known to be a response against stress induced-damage and often acts as a generic feedback to a variety of other equivalent stressors. However, such cyto-protective function in cancer may seem to be favourable for uncontrolled growth. This prompted us to analyse the expression of SatIII in human cancer cells. The present study reveals SatIII supports cancer cell survival, where it acts antagonist to the growth inhibitory drugs. Knockdown of SatIII restores functional p53 that in turn facilitates cancer cell death. Also, knockdown of SatIII is essential in sensitizing cells to chemo-treatments, hence, suggesting SatIII regulation in human malignancies facilitates an unknown mechanism of cellular protection against stress induced-death.

## 2. MATERIALS AND METHODS

### 2.1 Cell culture and treatments

The experiments were carried out using HeLa (cervical), HCT-15 (colon) and ZR-75-1 (breast) cancer cell lines obtained from the National Centre of Cell Science (NCCS), Pune, India. HeLa cells were grown in Dulbecco’s modified Eagle’s medium (DMEM, Sigma Aldrich^®^ Chemical Pvt. Ltd., New Delhi, India); HCT-15 and ZR-75-1 cells were grown in Roswell Park Memorial Institute medium (RPMI 1640, Gibco^®^, Thermo-Fisher Scientific, India), both supplemented with 10% fetal bovine serum (FBS) and 1% antibiotic (100 units/ml penicillin and 100mg/ml streptomycin). The culture conditions were maintained at 37°C and 5% CO_2_. The cells were grown in gelatin-coated sterile petridishes and sub-passaged as required for the experiments. The cells grown on 10mm 4 well-plates (on 0.1% gelatin-coated sterile coverslips) were allowed to float in a water bath maintained at 42°C for 1hr for heat shock (HS) treatment (Goenka et al., 2016). The cell confluency was maintained at 0.5×10^5^ prior to heat shock treatment. Another set of cells were left with no heat shock (NHS) at 37°C and harvested together with the heat shocked-cells.

### 2.2 Treatments with anti-cancer drugs

The cells were treated with 1μM Staurosporine, 0.2mM 5-Fluorouracil (5-FU) and 2.4μM of Cisplatin for 24h and processed for RNA FISH. One set of treated cells were given heat shock at 42°C for 1hr prior to *in situ* identification of SatIII transcripts, while the second set of cells were only drug-treated and left without heat shock prior to cell survival assay or RNA-FISH. Cell viability assay was done to determine the treatment concentration and duration as detailed in table 2 (supplementary SIII).

### 2.3 Knockdown approach and overexpression constructs

The cells were transfected with HSF1 knockdown construct (shRNA) purchased from Open Biosystem Co. and was validated in study by Sengupta et al. (2011). Transfection was achieved using TurboFect transfection reagent (Thermo Fisher Scientific^®^), for 36h at 37°C. The concentration of shRNAi-HSF1 was 100ng/μl. The HSF1 knockdown was coupled with the green fluorescent protein (GFP) expression construct (300ng/μl) for easy identification of transfected cells during fluorescence detection. The drug treated cells were transfected with either SatIII antisense or sense expression constructs with a desirable concentration of 250ng/μl (Table 3, supplementary SIII). The SatIII overexpression construct was made by subcloning of 158-bp fragment consisting of the SatIII repeat derived from 9q12 chromosome locus and cloned in pGEM2-98 bacterial vector into pcDNA3.1, the mammalian expression construct (pGEM2-98 was a kind gift from Dr. Caroline Jolly, INSERM, France; Goenka et al., 2016). The empty pGEM2-98 (250ng/μl) bacterial vector and pcDNA (200ng/μl) mammalian expression construct was used as controls. The p53 overexpression construct (400ng/μl) and shRNAi approach for p53 knockdown (400ng/μl) was used in couple with the green fluorescent protein (GFP) expression construct (300ng/μl). Transfection was achieved using TurboFect transfection reagent (Thermo Fisher Scientific^®^), for 36h at 37°C as per the manufacturer’s protocol.

### 2.4 Antibodies used for FISH and immunostaining

The following primary antibodies were used for the experiments: anti-digoxin (Jackson ImmunoResearch, USA; dilution 1:250), anti-HSF1 (Cell Signaling Technology^®^, USA; dilution 1:400), and anti-p53 (Cell signaling Technology^®^, USA; dilution 1:4000). The secondary antibodies used were: Streptavidin-Cy3 (1:2500), anti-Rabbit Alexa Fluor (1:500), anti-Mouse FITC (1:300) purchased from Jackson ImmunoResearch^®^, USA) as mentioned in Goenka et al. (2016).

### 2.5 SatIII probe synthesis

The *in vitro* synthesis of digoxigenin (DIG)-labeled antisense SatIII probe (158 bp SatIII repeat cloned in 1μg of PGEM-2-98 construct) with T7 RNA polymerase was carried out using the DIG labeling kit from Roche Products Pvt. Ltd, Mumbai, India (Goenka et al., 2016). The entire procedure was carried out in 37°C and the RNA was obtained overnight by 4M lithium chloride (LiCl) salt-precipitation in 1X TE. The RNA transcript was subjected to ethanol wash followed by air dry and dissolved in DEPC water. The product was further checked on 2% EtBr stained agarose gel and NanoDrop-quantified to achieve a desirable concentration of 250ng/μl (SpectraMax® M3 Multi-Mode microplate reader).

### 2.6 Fluorescence in situ hybridization (FISH) of SatIII transcripts

For the detection of SatIII transcripts, 5’-Dig-labeled complementary sequence to nucleotides 97–106 of the satellite III (5’ATTCCAATCCATGCCATTCC-3’) sequence was used (Sengupta et al., 2009). The cells were fixed with 4% paraformaldehyde (PFA) and processed for RNA *in situ* hybridization (RNA-FISH). The fixed cells were washed thoroughly in 1X PBS and permeabilized with 0.5% Triton X-100 on ice for 5mins. The DIG labeled-SatIII probe specific to chromosome 9q21 locus was then denatured with pre-hybridization buffer at 80°C for 5 min and re-natured on ice for 2min. The cells were further incubated overnight at 42°C with (DIG)-labeled antisense SatIII probe in hybridization buffer containing 1μg/ml yeast tRNA, 2X SSC, 50μg/ml heparin, 0.1% Triton X-100, 2% blocking agent and 25% formamide. The cells were then processed for the post-hybridization washes and subjected to the biotinylated-anti-DIG antibody (1:250) for 1 h at 37°C. The signals were amplified by streptavidin-Cy3 (1:2500) secondary antibody. The nuclei were stained with 2% DAPI and DABCO mounted after several washes with 1X PBS.

### 2.7 Cell viability assays

At the end of treatment, the cells were incubated with 0.5 mg/ml 3-(4,5-dimethylthiazol-2-yl)-2,5-diphenyltetrazolium bromide (MTT) for 4h at 37°C prior to harvesting. The purple formazan crystals formed at the end of incubation were dissolved in DMSO and the metabolic product was measured at 570 nm having a background reduction at 690nm by a colorimetric method using spectrophotometer (BioRad plate reader^®^ using SoftMax Pro6 2.2^®^ software) as reported earlier (Goenka et al., 2016; Upadhyay et al., 2016).

### 2.8 RNA isolation and RT-PCR

The transfection efficiency was confirmed by RT-PCR (Supplementary, Figure SII) and FISH. For this, the total RNA was extracted from the nuclear fraction using Trizol reagent (Invitrogen, USA) and cDNA synthesis was carried out with around 5μg of total RNA as reported in Goenka et al. (2016). The cDNA efficiency was further checked by RT-PCR with primers for the amplification of SatIII transcripts.

### 2.9 Signal detection and imaging

Signal detection and imaging done using an epifluorescence microscope (Axiovision) with the ApoTome module attached (Carl Zeiss, Bangalore, India). Images captured with 40X oil immersion lens using Zeiss Axio vision SE64 Rel. 4.9.1® software and edited with Adobe photoshop 7.0^®^. A minimum of 200 cells were accounted for blinded scoring (Nikon Eclipse 80i, Japan) and documented and graphically represented using Microsoft Excel 2010^®^.

### 2.10 Statistical analysis

Mean±s.d. was calculated for every experimental data and graphs were plotted with error bars. The statistical significance was analysed with unpaired Student’s *t*-test using Graph pad^®^, where a *P*≤0.05 was considered to be statistically significant.

## 3. RESULTS

### 3.1 Cancer cells HCT-15 and ZR-75-1 express SatIII independent of heat shock

We examined SatIII expression in vast majority of cancer cells (Table 1, Supplementary III) and we observed that colon and breast cancer cells expressed SatIII RNA independent of heat shock. This was an intriguing observation as both the cancer cells expressed SatIII under normal conditions. SatIII expression in human cancer cell lines, like HeLa and HEK is strictly heat regulated, where it facilitates cell-defense mechanism by activating the heat shock response pathway (Jolly et al., 2004; Sengupta et al., 2009). Under heat shock condition, majority of (>76%) HeLa cells were positive for SatIII-nSBs and consistent to earlier results, we here showed that the treatment with heat shock led to the formation of 5 to 6 nSBs per nucleus in majority of cells. Alternatively, we observed both HCT-15 (76%) and ZR-75-1 (69%) cells expressed SatIII RNA independent of heat shock in a significant proportion. This exceptional observation contradicted with the established notion of heat shock induced-SatIII expression, and therefore, we marked unstressed HeLa as negative control and heat shocked-Hela cells as positive control for our experimental cells (Figure 1a and Figure 1b). The quantitative analysis of SatIII RNA expression in HCT-15 and ZR-75-1 further revealed the number of SatIII-nSBs was two or more per cell. Heat shock acts as an additional stress to the cancer cells; therefore, SatIII-nSBs were detected considering heat shocked-HeLa as a positive control for the same. Elevation in temperature or heat shock in HCT-15 and ZR-75-1 enhanced SatIII expression and increased the number of nSBs per cell with large punctuated signals (Figure 1a) suggesting a hyperactive heat shock response consistent at the stages of tumorigenesis (Tilman et al., 2012). This raised the next question whether SatIII expression in cancer cells follows a similar mechanism to that of the expression of SatIII in heat shock conditions. To check this, we further looked into the expression of HSF1 and its colocalization with SatIII RNA in Hela, HCT-15 and ZR-75-1 cells.

**FIGURE 1.**
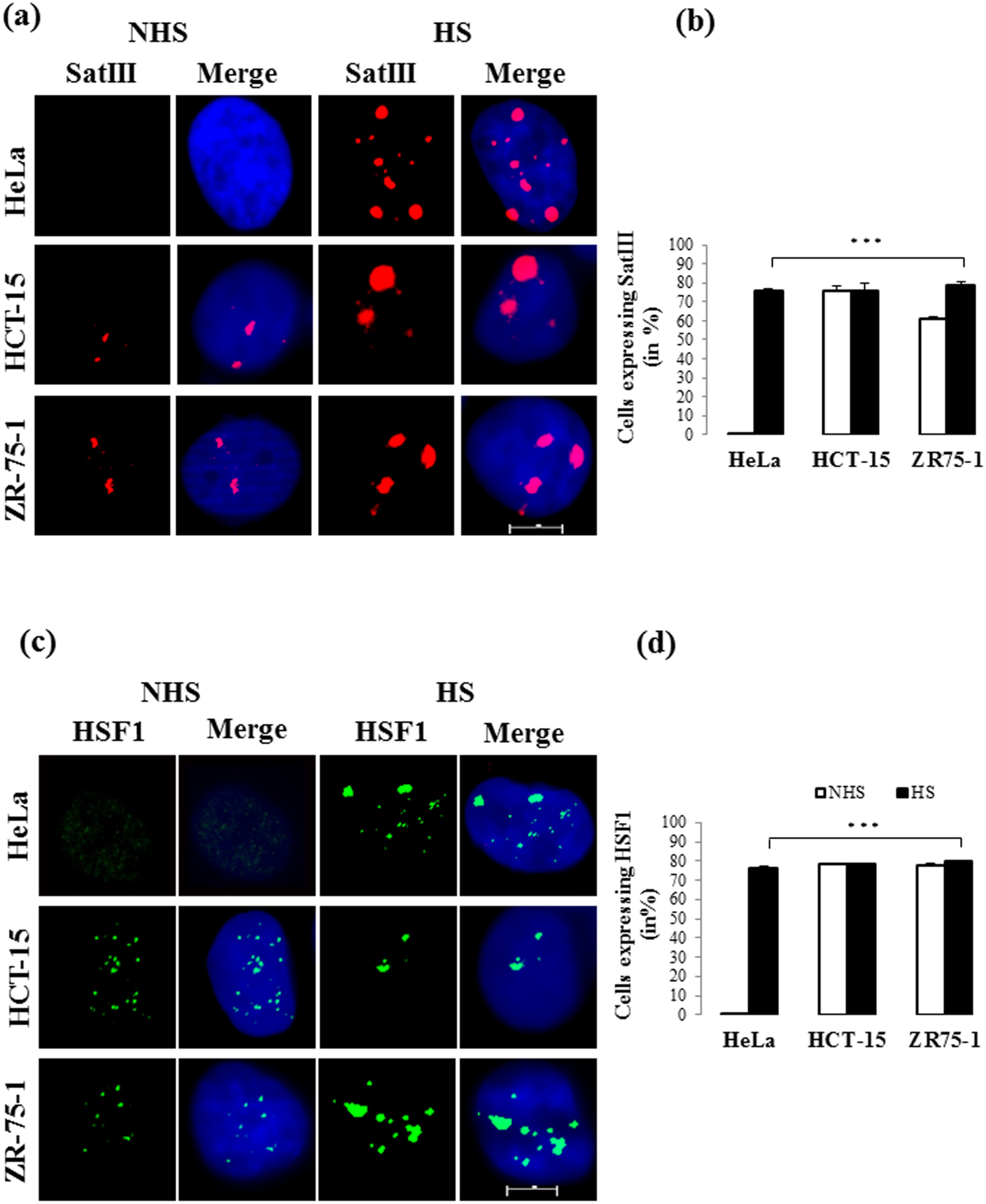
Cancer cells express SatIII and HSF1 independent of heat shock. (a) Immuno-FISH images showing SatIII positive nSBs in HCT-15 and ZR-75-1 nuclei are formed without heat shock; Thermal stress elevates the level of transcription (middle and lower panel of 1a). Note the expression of SatIII in HeLa is strictly heat shock dependent (upper panel). The proportion of cells with SatIII positive nuclei is presented in (b). (c) Constitutively active HSF1 in cancer cells require no heat shock for their expression in HCT-15 and ZR-75-1 nuclei. HSF1 expression in HeLa is tightly heat regulated as shown in the upper panel. The quantified data is presented in graph (d). NHS=no heat shock, HS=heat shock. Unpaired Student’s t-test, P *** ≤ 0.0005. Scale bar: 10μM

### 3.2 SatIII expression is independent of HSF1 in HCT-15 cells

The unstressed HeLa is known to have inactive HSF1 in the cytoplasm that translocates into the nucleus upon heat shock (Jolly et al., 2004; Sengupta et al., 2009). However, in HCT-15 and ZR-75-1 cells, we found HSF1 to form stress-specific signals in the non-heat shocked-cells similar to that of its expression in the heat shocked HeLa. Undoubtedly, treatment of cancer cells with heat shock led to the formation of large and intense HSF1 positive signals (Figure 1c and Figure 1d). We were further interested to see whether HSF1 and SatIII co-localizes in these two cancer cells. For the same we went for an endogenous HSF1 and SatIII signal detection which revealed that SatIII and HSF1 colocalizes in a very small proportion of cells, around 21% in both the colon and breast cancer lines. However, majority of cells, nearly 79% though positive with both the signals, showed distinctly punctated and non-colocalized SatIII and HSF1 in their nuclei. Heat shock increased the proportion of colocalized signals in the cells to 65% in HCT-15 and 53% in ZR-75-1, represented in Figure 2b, Figure 2f and Figure 2h. After this observation, it was essential to see whether HSF1 is really required by these two cancer cells for SatIII transcription. For this, we immediately knocked down HSF1 from HCT-15 and examined SatIII transcription. Surprisingly, HCT-15 continued to express SatIII-positive nSBs in a significant proportion of cells (76% and 79%) and showed no change in SatIII expression even after HSF1 was knocked down (Figure 2d and Figure 2g), both in control and heat shock conditions. HSF1 knock down in HeLa, however, resulted in complete absence of SatIII expression in the heat shocked-cells reinforcing earlier reports stating SatIII transcription in HeLa is strictly HSF1 dependent (Jolly et al. 2004; HSF1 RNAi approach by Sengupta et al. 2009). Thus, SatIII transcription in cancer cells appears to be driven by oncogenic stress and may involve transcription factors other than HSF1. The ZR-75-1 cells were too sensitive to HSF1 knockdown; therefore for this cell line we limited our results till the colocalization study.

**FIGURE 2.**
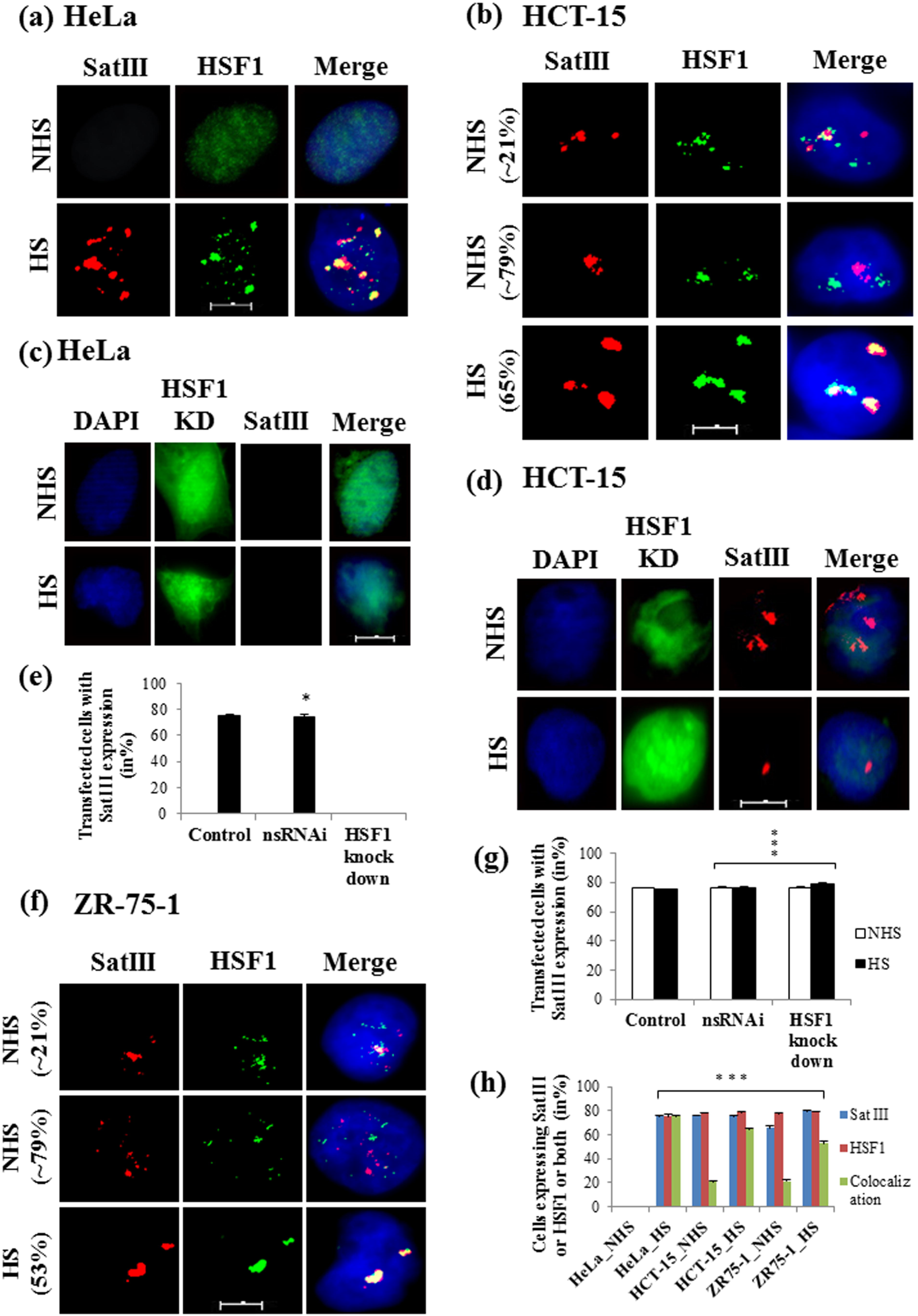
Cancer cells express SatIII transcripts independent of HSF1. (a) Immuno-FISH images showing SatIII and HSF1 undergoes complete colocalization (yellow puncta) in heat shocked-HeLa. (c) HSF1 is essential for SatIII transcription in HeLa; note knockdown of HSF1 with shRNA approach did not express SatIII even after heat shock. The quantified data is presented in graph (e). (b) SatIII and HSF1 showing partial colocalization in HCT-15 (upper panel), where the cells without colocalized nSBs were detected with closely located SatIII and HSF1 signals. Heat shock increases the % of co-localization of HSF1 and Sat III as seen in lower pane; 1b (d) HSF1 knockdown did not affect SatIII transcription in HCT-15 cells, rather resulted in SatIII associated nSBs in both the non-heat shocked and heat shocked-cells. The percentage of SatIII expressing cells in HSF1 knocked down HCT-15 is presented in graph (g). (f) SatIII and HSF1 showing partial colocalization in ZR-75-1 cells (upper panel). Heat shock increases the proportion of cells with co-localized puncta **(lower panel 1f)**. The quantified colocalization data for all the three cell lines are presented with bar diagrams in (h). NHS=no heat shock, HS=heat shock. Unpaired Student’s *t*-test, P * ≤ 0.05, *** ≤ 0.0005. Scale bar: 10μM

### 3.3 SatIII transcription promotes proliferation of HCT-15 (colon) and ZR-75-1 (breast cancer) cells

Several studies have emphasized on the transcriptional and epigenetic status of the pericentromeric satellite transcripts that if altered enhances the risk of malignancies (Zhu et al., 2018; Giordano et al., 2020). Since, we observed that the level of SatIII expression varies between unstressed and heat shocked-human colon and breast cancer cells, we further decided to investigate whether SatIII transcripts are required for cancer cell progression. For this, we primarily knocked down SatIII in HeLa cells. When compared to control set (non-transfected, or transfected with an empty vector pcDNA or pGEM2-98), knockdown of SatIII in the heat shocked-HeLa led to significant cell death, around 42% as represented with bars in Figure 3a. However, loss of SatIII in unstressed HeLa did not affect the rate of cell survival, hence, suggested SatIII transcripts are essential for the survival of HeLa cells only under thermal stress condition. Overexpression of SatIII transcription in unstressed HeLa reduced cell survival from 100% to 83%, which was further decreased to 66% upon treatment with heat shock (Figure 3a). This indicated a toxic effect of SatIII transcripts when aberrantly expressed in the heat shocked-cells as already established in an earlier study by Goenka et al. (2016). With this reference, we then knocked down SatIII in unstressed HCT-15. As compared to control with 100% survival rate, SatIII knockdown significantly lowered the proportion of viable cells to 54% (Figure 3b). Treatment with heat shock even reduced the percentage of surviving cells to 41% suggesting a failure of its heat shock response-assisted cyto-protective role when depleted from the proliferating cells. The SatIII transcripts when overexpressed in the HCT-15 cells, increased cell survival to 93% as compared to the 100% control cells or even SatIII knocked down cells (54%) (Figure 3b). Heat shock to HCT-15 cells in SatIII overexpressed condition, however, reduced the rate of cell viability to 87% (from 100%), suggesting an aberrant SatIII expression may be associated with cytotoxicity. Next, we knocked down SatIII from ZR-75-1 cells and examined cell viability. There was indeed a fall in cell survival rate to 61% (from 100%) in the non-heat shocked-cells (Figure 3c). Treatment with heat shock, however, contradicted with our earlier observations and increased the proportion of viable cells to 82% when compared to the control, heat shocked or SatIII depleted HeLa and HCT-15 cells (Figure 3c). Interestingly, overexpression of SatIII in ZR-75-1 cells further accelerated cell growth to 82% in control conditions and achieved maximum viability of 92% when given a heat shock treatment. This might indicate heat shock triggered SatIII to increase the rate of cell survival to significant fold (Figure 3c). Thus the overall observation suggests SatIII transcription in certain human cancers favours cell survival. Heat shock either increases the load of cytotoxicity and lowers cell survival rate, or, potentiates SatIII to continue its cell survival function depending upon the type of cells or inner cellular milieu. As loss of SatIII transcripts majorly affected cell viability, hence SatIII lncRNA may play crucial role in cancer cell death.

**FIGURE 3.**
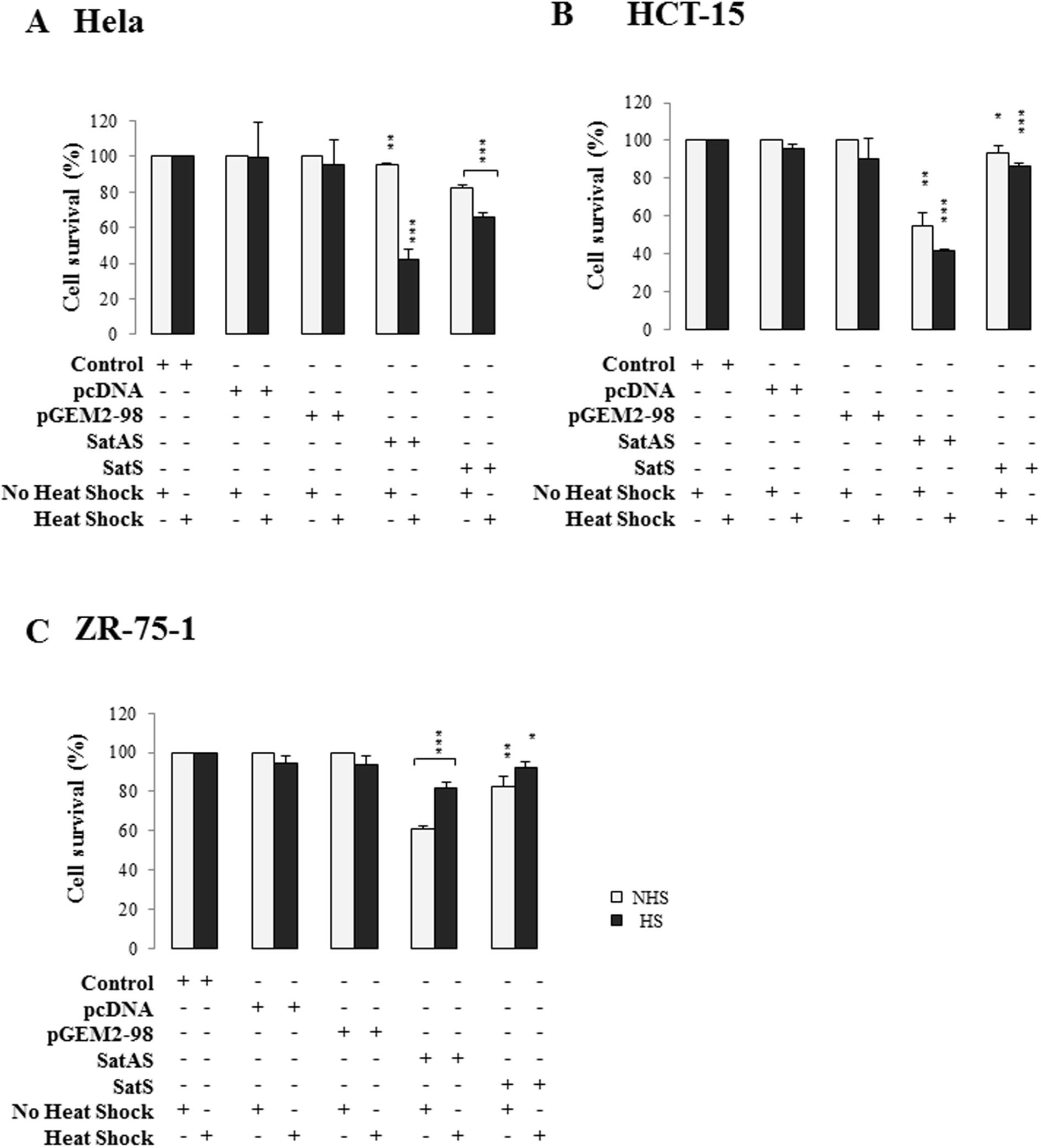
SatIII facilitates survival of colon and breast cancer cells. MTT analysis revealed a significant proportion of cells undergoes apoptosis when SatIII is knocked down in HCT-15 (b) and ZR-75-1 (c) lines. Note the unstressed HeLa cells (a) continue to survive as they do not express SatIII without heat shock treatment. In contrast, SatIII overexpression led to cancer cell survival as represented in graphs (b) and (c). pcDNA and pGEM2-98 were empty vectors and considered as controls. SatS=SatIII overexpression, SatAS=SatIII knockdown constructs. NHS=no heat shock, HS=heat shock. Unpaired Student’s *t*-test, P * ≤ 0.05, ** ≤ 0.005, *** ≤ 0.0005

### 3.4 Chemotherapeutic drugs induce expression of Satellite III RNA in Hela cells

As seen in the previous section, SatIII possesses a unique feature of cyto-protection when overexpressed in the colon and breast cancer cells. This tempted us to further analyse whether SatIII continues to protect cancer cells when treated with growth inhibitory drugs. Since HeLa induces SatIII only when treated with heat shock, the effect of anti-cancer agents (supplementary, Figure SI) on SatIII induction was primarily tested on HeLa and the *in situ* data were recorded for further reference. HeLa being a control cell line, was tested with all the three growth inhibitory drugs: staurosporine, an apoptosis inducer; 5-FU, a chemo-drug specific to colon cancer cells, and cisplatin, a chemo-drug specific to the breast cancer cells. Intriguingly, all the three anti-neoplastic drugs induced SatIII in HeLa even without heat shock in a significant proportion of cells. Staurosporine induced SatIII expression in 76% of HeLa cells while 90% of the cells expressed SatIII when treated with cisplatin. Treatment with 5-FU induced SatIII in maximum number of cells; around 94% of HeLa were detected with SatIII positive nuclei as compared to the non-treated control cells which did not express SatIII until given a heat shock treatment (Figure 4a and Figure 4a1). The brightest signals formed during heat shock treatment suggest an increased amount of target RNA in the SatIII transcriptional loci (Figure 4a). Overall a distinct and reproducible pattern was observed with regard to the number of nSB positive cells, and for the number of nSBs observed in each cell for each of the three stressors tested. Therefore it confirmed that the chemotherapeutic drugs used in our study induced SatIII expression in Hela cells.

**FIGURE 4.**
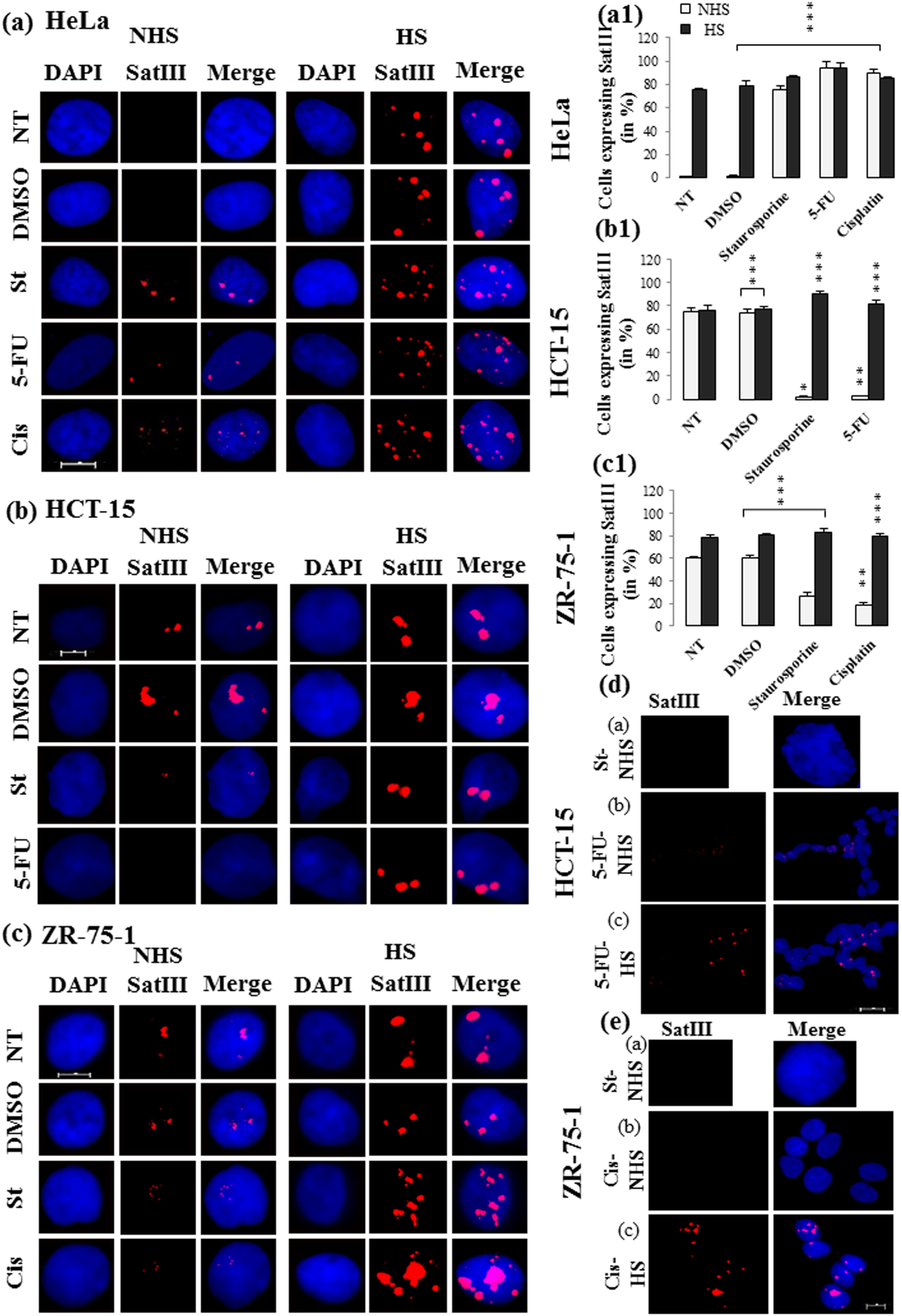
Anti-cancer drugs reduce SatIII transcription and facilitate cancer cell death. The extent of SatIII expression was observed in all the three cell lines after treatment with 1μM staurosporine, 0.2mM 5-FU and 2.4μM cisplatin for 24h. (a) HeLa shows drug induced-SatIII transcription in non-heat shocked cells. Heat shock combined with the cytotoxic effect exerted by chemo-drugs and together increased the level of SatIII in HeLa. The percentage of drug treated HeLa cells harbouring SatIII positive nSBs are presented in (a1). (b) Staurosporine and 5-FU effectively reduced SatIII transcription in HCT-15 cell. SatIII expression showed an aberrant increase upon heat shock. The quantified cells with SatIII positive nSBs per treatment is presented in graph (b1). (c) Staurosporine and cisplatin inhibited SatIII transcription in unstressed ZR-75-1 as compared to the control set of cells. In contrast, heat shock led to aberrant increases in SatIII transcription. The proportion of SatIII positive cells upon treatments are quantified and presented in (c1). (d) Staurosporine inducing condensation of nucleus is shown in the DAPI stained nucleus of HCT-15 cell (panel a). Treatment with 5-FU led to morphologically healthy cells and showed no SatIII transcription (panel b). The cells continued to grow in clumps when given a heat shock which also enhanced SatIII transcription (panel c). (e) Staurosporine induced nuclear disintegration as seen in the DAPI stained apoptotic ZR-75-1 cells (panel a). Cisplatin released cell clumps with reduction in the expression of SatIII RNA (panel b). The cells revived its malignant phenotype (cell clumping) when given a heat shock with enhanced SatIII transcription (panel c). NT= Non treated, St= Staurosporine, 5-FU= 5-Fluorouracil, Cis=Cisplatin. DMSO was used as a vehicle. NHS=no heat shock, HS=heat shock. Unpaired Student’s *t*-test, P * ≤ 0.05, ** ≤ 0.005, *** ≤ 0.0005. Scale bar: 10μM

### 3.5 Chemotherapeutic drugs suppress SatIII expression in colon cancer cells

In the next step, we treated the colon cancer cell line, HCT-15 with staurosporine and 5-FU, and observed their effects. Staurosporine is known to modulate the lipid constitute while inducing apoptosis in the colon cancer cells (Solar et al., 2015). Staurosporine treatment in HCT-15 led to only 2% SatIII positive cells as compared to the control group that induced SatIII in about 76% of the non-treated and non-heat shocked HCT-15 cells (Figure 4b and Figure 4b1). Exposure of staurosporine treated HCT-15 cells to heat stress increased the proportion of cells expressing SatIII to 91% (Figure 4b). We also stained the treated cell nuclei with DAPI and observed under fluorescence microscope. The cells revealed nuclear shrinkage (panel a, Figure 4d), corroborating earlier report that suggests staurosporine alters cellular morphology by inducing nuclear condensation and forming electron dense bodies during the stages of apoptosis (McKeague et al., 2003). This may indicate an inhibitory effect of staurosporine on SatIII expression in order to make HCT-15 undergo cell death.

It is interesting to know that the colon cancer cells exhibit N-glycans on their surface that if altered leads to malignant cell fate (Gao et al., 2014). 5-FU specifically targets colon cancer cells, interferes with their glycosylation and restricts cell growth. We therefore treated HCT-15 cells with 5-FU and observed its effect on SatIII expression. Intriguingly, 5-FU treatment resulted in complete absence of SatIII RNA in majority of the cells (Figure 4b). We detected null to around 2% of the HCT-15 cells with SatIII positive nuclei as compared to, non-treated group that usually exhibit around 76% of SatIII expressing cells (Figure 4b). Heat shock treatment increased the cumulative cell count of SatIII positive signals to 82% in compare to the unstressed cells (2%). The size of punctated SatIII loci was relatively large in heat shocked-cells, where three or more loci per cell transcribed SatIII than a null expression observed in the unstressed condition. We further checked for any variation in the nuclear phenotype arises due to growth-inhibitory drugs followed by a heat shock therapy. This led to two important observations; first, treatment with anti-proliferative drugs released cell clumps that otherwise exhibits a stringent adherence. Since chemo-drugs induced malignant cell death, we observed a reduced expression of SatIII RNA in the still-alive cells (panel b, Figure 4d). Second, we observed treatment with heat shock in the chemo-treated cells led to distorted nuclear phenotype with aberrant SatIII signals in the clustered cells; therefore, suggested a revival of their malignant state (panel c, Figure 4d).

### 3.6 Chemotherapeutic drugs suppress SatIII expression in breast cancer cells

Staurosporine is known to release mitochondrial cytochrome c and trigger caspase-dependent apoptosis in breast cancer cells (Xue et al., 2003). However, it may also act as a chemo-protective agent and arrest normal cells while allowing the tumor ones to undergo chemo-induced death (Murray et al., 2013). Since SatIII promotes malignant cell growth, we wanted to check whether staurosporine regulates SatIII expression while triggering apoptosis in breast cancer cells. For this, we treated ZR-75-1 cells with staurosporine and subjected them to *in situ* hybridization analysis for the detection of SatIII lncRNA. It was intriguing to observe that, treatment with staurosporine led to loss of SatIII transcripts in unstressed ZR-75-1 cells. Up to 27% of ZR-75-1 cells didn’t have SatIII RNA. This was in comparison to the control group (untreated) that usually expresses about 61% SatIII positive cells under normal condition (non-treated, no heat shock, Figure 4c and Figure 4c1). Treatment with heat shock however increased the proportion of cells expressing SatIII RNA to 84%. The proportion was higher than the untreated group that under heat shock expressed SatIII in 79% cells. Also the percentage was significantly higher than the 27% SatIII positive cells of the treatment group. We next observed the DAPI stained nuclei under fluorescence microscope and found staurosporine treated cells with condensed nuclear phenotype (panel a, Figure 4e).

Since cisplatin targets the breast cancer cells through multiple signaling pathways and regulates processes like replication and transcription to restrict their growth (Jiang et al., 2017), we next checked the effect of cisplatin on SatIII transcription in ZR-75-1 cells. We observed cisplatin exerted an inhibitory effect on breast cancer cells, and reduced SatIII expression in these cells. Only a small proportion of around 18% of ZR-75-1 cells showed SatIII accumulation (as compared to 61% untreated ZR cells), however, the signal intensity was somewhat undetectable (left bottom panel, Figure 4c). Elevation in temperature dramatically increased the proportion of SatIII expressing cells to 80% with remarkably large and aberrant SatIII expressing loci (Figure 4c). This might suggest heat stress in combination with cytotoxic drugs, led to an enhanced level of SatIII transcription in response to the cumulative stress. Alternatively, SatIII may have distinct pathways and roles to play in stress response and cancer. When analysed for morphological changes, the cisplatin treated cells looked phenotypically stable with no SatIII expression, suggesting elimination of dead cells from the tumor niche (panel b, Figure 4e). Treatment with heat shock, however, showed a speedy recovery from chemo-induced damage and restored malignant phenotype to the cells found positive with large SatIII transcribing loci (panel c, Figure 4e).

Therefore, the overall observation revealed that chemotherapeutic drugs reduced SatIII expression in the non-heat shocked-HCT-15 and ZR-75-1 cells. In contrast, the chemicals effectively induced SatIII in the unstressed HeLa similarly as found in the heat shocked cells. This suggests SatIII expression in cancer has distinct and specific role rather than a mere stress indicator. Most importantly SatIII can be transcriptionally repressed by the anti-cancer drugs for triggering cancer cell death and this can be a therapeutic intervention for future cancer treatments.

### 3.7 SatIII transcripts protect HCT-15 and ZR-75-1 cells against chemo drug-induced cell death

SatIII expression was repressed by specific chemotherapeutic drugs and as SatIII also has a role in cell survival, hence it was very exciting to see the role of SatIII in context of cell survival in chemo drug treated conditions. For this, we either overexpressed or knocked down SatIII in the drug-treated cell lines, and examined the extent of cell death.

The MTT results revealed treatment with 5-FU led to a significant percentage of cell death in HCT-15 cells, leaving around ∼62% viable cells in the culture as compared to 100% survival rate in non-treated control group (Figure 5a). Similarly, we observed that ZR-75-1 cells are sensitive to cisplatin treatment. Only a small proportion, about ∼15% cells survived as compared to the 100% viable cells from the non-treated group (Figure 5a). Therefore, after comparing the non-treated (control) and treated group, we here show the specific growth-inhibitory drugs prevent proliferation of the colon and breast cancer cells.

**FIGURE 5.**
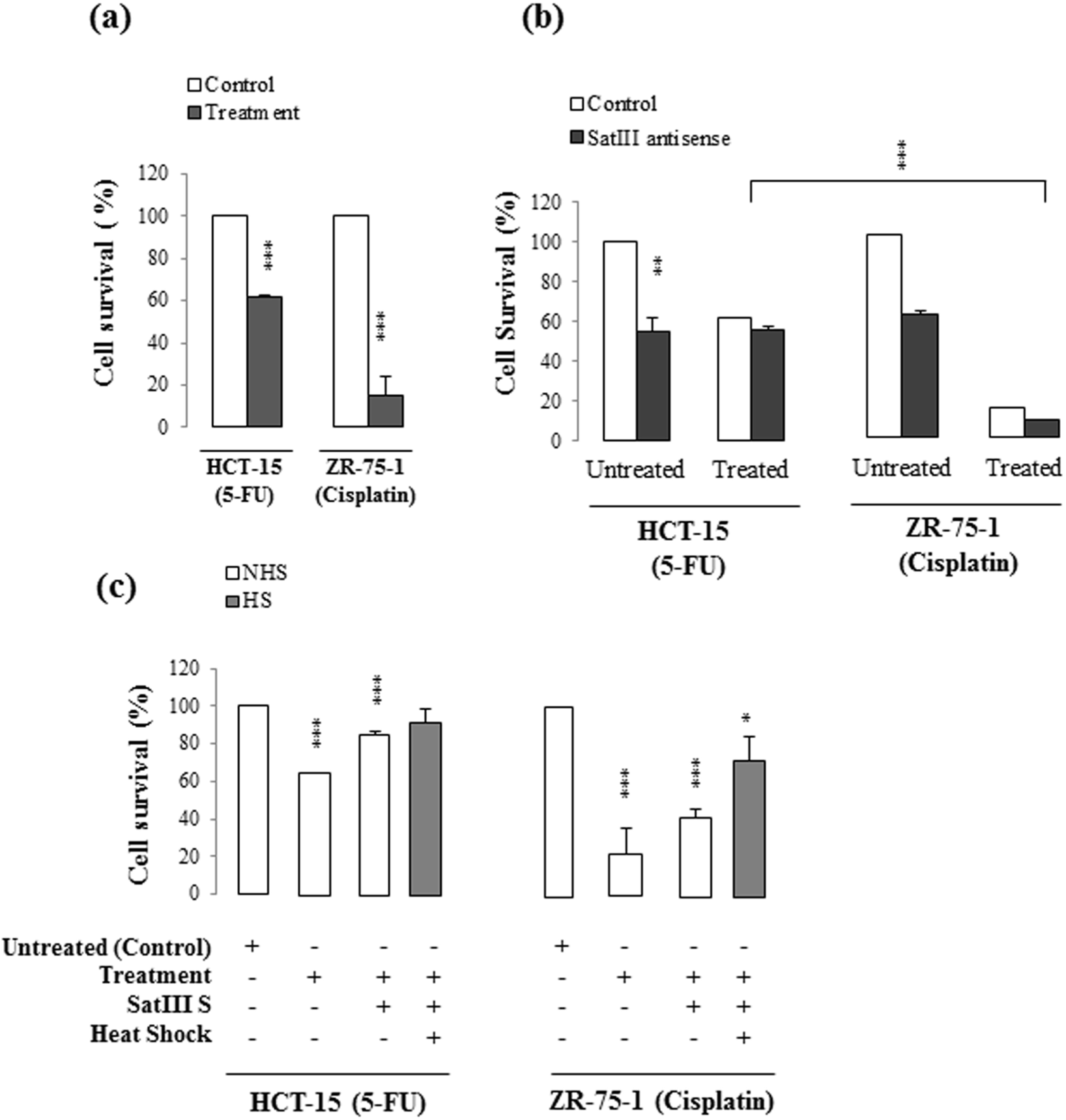
Overexpression of SatIII confers protection against chemo-induced cancer cell death. (a) Cell viability assay revealed 5-FU and cisplatin efficiently restricts survival of HCT-15 and ZR-75-1 cells respectively. The proportion of viable cells reduces with treatments in compare to the untreated control group. (b) Bar diagram represents knockdown of SatIII in drug-treated group induces cell death. The proportion of viable cells reduces than treatment group alone. The treatment group represents HCT-15 and ZR-75-1 cells that were treated with 5-FU and cisplatin respectively. (c) Overexpression of SatIII in both cells induced cell survival in compare to the SatIII knockdown group presented in graph (b). The proportion of viable cells was higher upon SatIII overexpression even after treatment with chemo-drugs. Heat shock in SatIII overexpressed group continued to facilitate cell survival. NHS=no heat shock, HS=heat shock. Unpaired Student’s t-test, P * ≤ 0.05, ** ≤ 0.005, *** ≤ 0.0005

To examine the effect of SatIII knockdown on the sensitivity of colon cancer cells to 5-FU, we transfected HCT-15 with SatIII antisense construct and exposed the cells to 5-FU at concentration 0.2mM. We then compared the treated and untreated group of cells that were transiently transfected with SatIII antisense construct. As shown in Figure 5b, SatIII knockdown in cells that were treated with 5-FU rapidly induced cell death to a significant proportion leaving only 55% viable cells. This was lower as compared to the cells that were received 5-FU treatment alone (Figure 5b, treated control, ∼62% cell viability) and was not transfected with SatIII antisense construct. The results therefore suggest SatIII knockdown together with chemo-drug, 5-FU increases the overall percentage of cell death. However, the rate of cell survival was similar to that of the untreated, SatIII knocked down group (∼54% cell viability). Thus, as shown in the bar diagrams, in both cases, (treated and untreated), knockdown of SatIII led to an increased proportion of cell death. Our results therefore suggest loss of SatIII transcripts sensitizes the colon cancer cells to 5-FU treatment and together leads to maximum cell death (Figure 5b).

We next treated the ZR-75-1 cells with 2.4μM cisplatin that involve DNA-DNA or DNA-protein interstrand crosslinks to induce apoptosis in breast cancer cells (Sigurðsson et al., 2014). Intriguingly, we here show knockdown of SatIII greatly impacts survival of ZR-75-1 cells. Loss of SatIII transcripts in cisplatin treated group led to only 8% cell survival as compared to the cells that received cisplatin treatment alone (Figure 5b, ∼15% cell viability). The results therefore suggest SatIII knockdown together with cisplatin treatment increases the overall percentage of cell death. Intriguingly, while comparing the untreated and treated groups, the proportion of cell death appears to be higher with SatIII knockdown in the chemo-treated group (8% cell viability). This was in comparison to only SatIII knocked down cells in the untreated group (∼61% cell viability). The overall observation therefore suggests SatIII confers a pro-cell survival function in response to cancer cell growth. Loss of SatIII in combination with apoptosis-inducing drug enhances cell death which can be very useful in cancer treatment.

We next wanted to check whether overexpression of SatIII transcripts in drug-treated condition would protect cancer cells from cell death. In HCT-15, overexpression of SatIII transcripts led to 85% cell survival as compared to the 100% cells of untreated control group (Figure 5c). The percentage of cells survived upon SatIII overexpression appears to be higher than the non-transformed cells that received 5-FU treatment alone (∼62%). Overexpression of SatIII in ZR-75-1 led to 42% cell survival as compared to 100% viable cells in untreated control group. However, the percentage of cell survival was higher in SatIII overexpressed group as compared to the non-transfected cells that received cisplatin treatment alone (∼15%). The results, therefore suggests, SatIII overexpression induces cell survival even after treatment with anti-cancer drug. In addition to this, overexpression of SatIII transcripts induces survival of both HCT-15 and ZR-75-1 cells (Figure 5c) as compared to SatIII knocked down group (Figure 5b). Hence, comparing all three conditions (treatment alone, treatment + SatIII AS, and treatment + SatIII OS), we suggest overexpression of SatIII gradually increases the rate of cell survival. This therefore suggests overexpression of SatIII protects cells from chemo-induced death. Heat shock in SatIII overexpressed group continued to facilitate cell survival (Figure 5c). This in HCT-15 led to 92% cell viability even after treatment with growth inhibitory drug. The same for ZR-75-1 cells led to 73% cell survival indicating heat shock potentiates SatIII to continue its cell survival function (Figure 5c). This correlates to our *in situ* data and corroborates to an elementary observation of heat shock mediated chemo-resistance reported earlier in several other studies (Arya et al., 2007; Matsunaga et al., 2014). There seems to be a possibility of cell toxicity due to prolonged overexpression of SatIII and hence a proportion of cells underwent apoptosis. This corresponds to the observation suggesting chronic heat shock condition can induce cell death (Stankiewicz et al., 2009). Our overall results therefore suggest SatIII overexpression responds similarly to heat stress as well as facilitates proliferation of colon and breast cancer cells.

### 3.8 SatIII knockdown leads to higher expression of p53 protein thereby triggering cancer cell death

Our previous *in silico* study showed a strong SatIII RNA and p53 protein interaction (Fig. 7 a and b). The SatIII-interacting residues present on p53 DNA binding domain appeared functionally significant in cellular stress and cancer, which, if altered, may lead to malignant cell fate (Chatterjee et al., 2019). Several research groups also have shown that p53 intrinsically interacts with specific lncRNAs in cancer (Chaudhary & Lal, 2017; Lin et al., 2019). To understand the mechanistic role of SatIII transcripts in cancer cell death, we further looked for any physical interaction of SatIII and p53. This was checked in Hela and also in our experimental cancer cell line, HCT-15. In HeLa, under the condition of heat shock, about 18% of the SatIII positive cells co-stained with p53 protein (Figure 6a). In contrast, we show the recruitment of p53 onto SatIII positive nSBs (18%) without a heat shock treatment in the HCT-15 cells (Figure 6b). Considering heat shocked-HeLa cells a positive control, an additional heat shock treatment was given to the HCT-15 cells. In this, p53 protein now co-stained with about 16% of the SatIII positive nuclei. The large and bright colocalized signals in Figure 6b revealed that the cancer cells partially recruit p53 onto SatIII-associated nuclear stress bodies.

**FIGURE 6.**
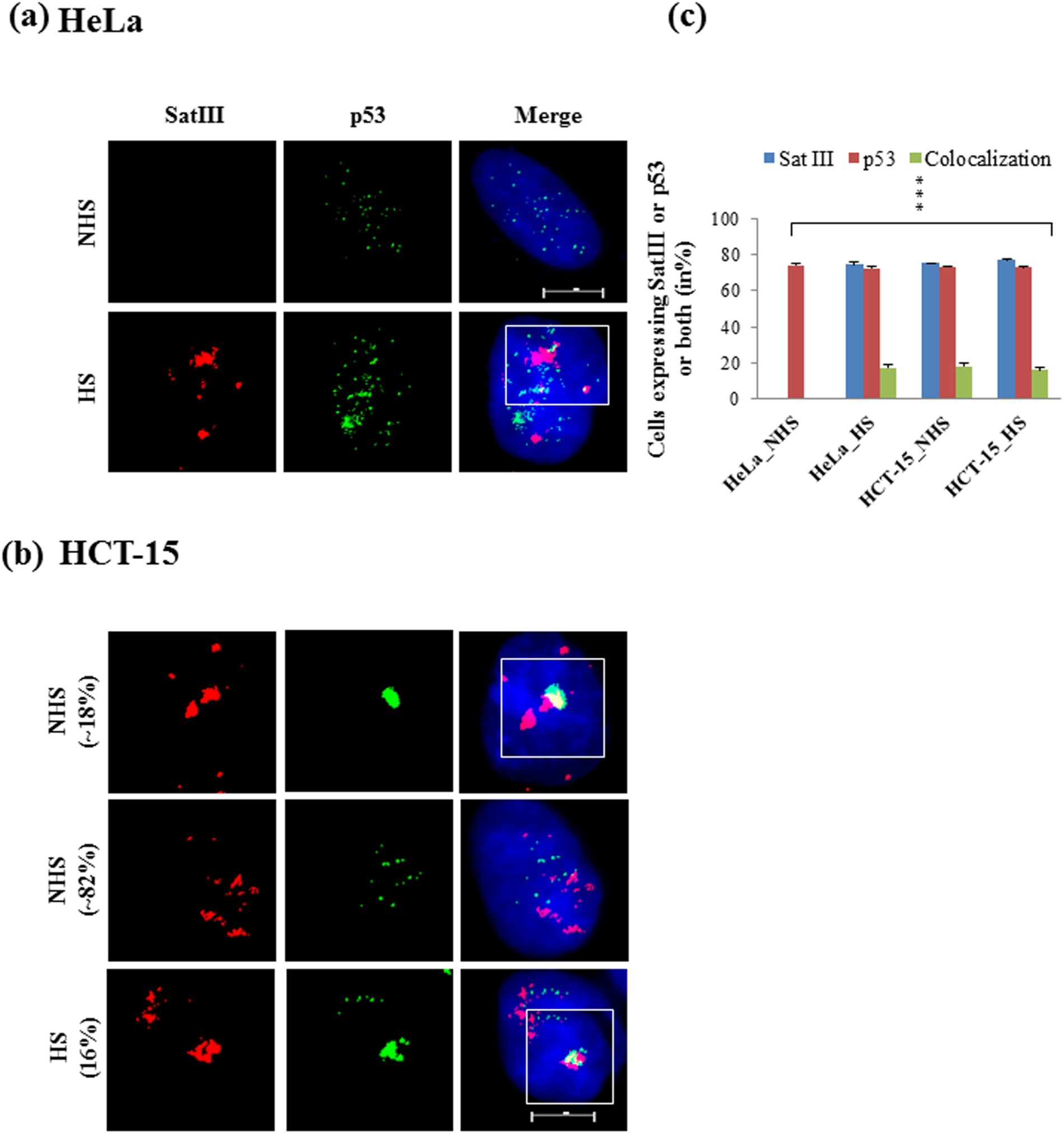
Cancer cells harbour partially colocalized SatIII and p53 signals onto nuclear stress bodies. (a) Immunostaining image showing in HeLa, under condition of heat shock, p53 partially colocalizes with SatIII positive nSBs. Note the minimal expression of p53 in HeLa (upper panel). The control cells did not express SatIII. (b) In HCT-15, the SatIII positive nuclei were co-stained with p53 protein in both unstressed and heat shock conditions. The colocalized puncta (yellow) in HCT-15 are brightest suggesting an increased amount of target RNA and protein in the cells. The quantified data for SatIII and p53 colocalization is presented with bar diagrams in (c). Squares in white identify the SatIII positive nSBs that also co-stained for p53 protein. NHS=no heat shock, HS=heat shock. Unpaired Student’s t-test, P *** ≤ 0.0005. Scale bar: 10μM.

**FIGURE 7.**
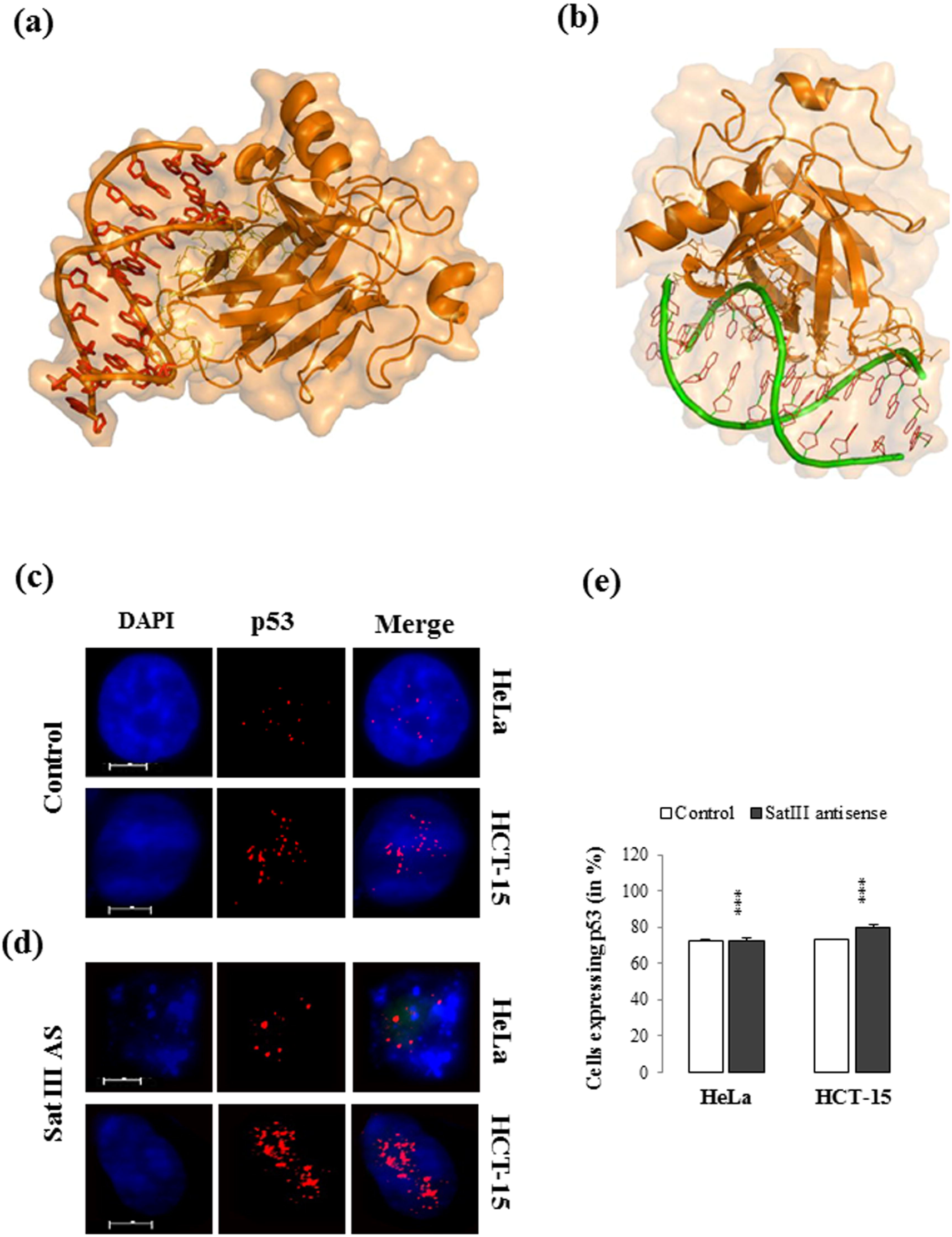
SatIII knockdown induces p53 expression in cancer cells. (a) Docked complex representing interactions between SatIII and p53 with interacting amino acid residues present on p53 DNA binding domain (yellow) (adapted from Chatterjee et al. 2019). (b) Molecular interaction analysis by PatchDock beta v1.3 shows interaction between mutp53 and SatIII leads to change in amino acid residues present on p53 DNA binding domain (pink) (adapted from Chatterjee et al. 2019) (c) In control condition (unaltered Sat III levels), p53 expression is low in HeLa cells (upper panel). In HCT-15, in control condition, 2 to 3 p53 puncta along with small dispersed granules per cell is found (lower panel). (d) SatIII AS represents set of cells in which SatIII has been knocked down with SatIII antisense oligo. In HeLa, SatIII knockdown induces p53 expression and stress-specific signals are formed. Nuclear fragmentation is seen in HeLa cells in SatIII knockdown condition (upper panel). In HCT-15, in SatIII knockdown cells, p53 expression was stronger and spread over the nuclei. (e) Bar diagrams representing the number of cells expressing p53 in control and in SatIII knockdown conditions. This is not related to change in p53 protein expression level in SatIII knocked down cells. Both the control and SatIII AS are heat shocked cells. SatIII AS=SatIII antisense construct. The SatIII antisense oligo was used in couple with the green fluorescent protein (GFP) expression construct for easy identification of transfected cells. Unpaired Student’s t-test, P *** ≤ 0.0005. Scale bar: 10μM.

In unstressed condition, the rapid turnover of p53 maintains the protein at a low level for normal cell functioning (Midgley & Lane, 1997). However, the level of p53 expression increases with stress, and the active p53 protein is translocated into the cell nucleus (Ho et al., 2019). To investigate further into SatIII and p53 interactions, we knocked down SatIII in both HeLa and HCT-15 cells. Since SatIII expression in HeLa cells is heat shock dependent, the entire set of SatIII knockdown experiment was performed under the condition of heat shock, for both HeLa and HCT-15 cells. Heat stress stabilizes p53 (Miyakoda et al., 2002), though we showed the protein expression was low in the nuclei of HeLa cells (Figure 7a, upper panel). In HeLa, upon SatIII knockdown, there was an increased expression of p53 protein. The brighter signals along with the large number of punctated p53 loci was surprising and was present in about 72% of the HeLa cells, suggesting an enhanced expression of target protein (Figure 7a, lower panel). The apoptotic nuclei showed stress-specific p53 signals that greatly differed from its low level expression in unstressed HeLa cells (Figure 7c, control panel).

In HCT-15 cells, heat shock led to the formation of 2 or more stress specific puncta (in ∼73% of cells) along with small dispersed granules per cell nuclei (Figure 7b, upper panel). The stress-specific expression of p53 suggests that in cell culture, tumor-associated stress is a strong inducer of p53 and formation of nuclear stress bodies. In cancer cells, p53 aggregations can dysregulate the proteostatic network. This possibly includes a hyper-active, oncogenic heat shock response to which the tumors get addicted (Smet al., 2017). We observed that loss of SatIII transcripts increased the availability of p53 protein in HCT-15 cells (80%). The level of p53 aggregation in HCT-15 was much higher than its minimal expression in the non-transformed cells. The activated p53 then induces cell death as shown in the DAPI stained apoptotic nuclei (Figure 7b). The proportion of HeLa and HCT-15 cells expressing p53 upon SatIII knockdown is presented in graphs (Figure 7c). This is not related to the change in p53 protein expression level in SatIII knocked down cells. Taken together our results suggest that while SatIII knockdown enhances the level of p53, its transcriptional status then greatly impact the survival of cancer cells.

## 4. DISCUSSION

The regulatory function of non-coding RNAs in fundamental cellular processes has been detected over the years. In particular, the involvement of long non-coding RNAs (lncRNAs) with nuclear organization and chromatin remodelling often makes them responsible for growing risk of human malignancies (Bartonicek et al., 2016; Sanchez Calle et al., 2018; Chatterjee & Sengupta, 2019). We here provide a novel insight into the regulatory role of human satellite III (SatIII) lncRNAs in colon and breast cancers. We show SatIII transcription is a response to oncogenic stress. This suggests SatIII is active during many physiological processes in addition to its heat induced-expression and accumulation onto nuclear stress bodies (Jolly et al., 2002; Valgardsdottir et al., 2008; Sengupta et al., 2009; Goenka et al., 2016). Cancer cells harbour ever-active HSF1 that forms stress-specific granules corroborating the notion that the constitutive activation of HSF1 is essential for the stages of tumorigenesis (Dai & Sampson, 2016). The ∼20% proportion of SatIII-nSBs still colocalizes with HSF1 in the colon and breast cancer cells suggests HSF1 potentiates lncRNAs to drive tumor cell growth. This appeared to be similar with the report HSF1 empowers NEAT1 to confer oncogenic activity (Watanabe et al., 2018). An HSF1 independent SatIII transcription however indicates involvement of other transcriptional factors (Valgardsdottir et al., 2008; Sengupta et al., 2009), or tumor specific kinases for the transcription of SatIII loci in human cancer cells. We therefore propose induction of SatIII lncRNA in cancer cells is an exception of its expression in the conventional heat shock response pathway. It has been known for a while now that the lncRNAs either bear oncogenic potential, or act as tumor suppressors depending upon their cell or tissue specific expressions (Beckedorff et al., 2013). We suggest SatIII favours a persisting malignant growth, which when knocked down leads to a significant proportion of cell death. Notable, the overexpressing satellite RNAs become sensitive to DNA replication stress and induce malignant lesion (Zhu et al., 2018). Although our study is first of its kind to comment on the expression of SatIII in cancer, however, a line of evidence suggests a bromodomain protein, BRD4 interacts with HSF1 in the nSBs and upregulates SatIII in stress induced neoplasm (Hussong et al., 2017). The exponential growth achieved by the unstressed cells suggests forced SatIII transcription may mimic the HSR mechanism and trigger cell survival in human even in no heat shock condition. Therefore, SatIII exhibits a multifaceted role through its involvement in both the regulation of heat shock response and cancer, two distinct yet inter-regulatory pathways that have huge impact on the overall physiological outcome of stress. The principle of chemo-treatment includes the limited DNA damage repair ability of tumor cells that makes them susceptible to radiation than normal cells. Growth preventing drugs like staurosporine, 5-FU, and cisplatin majorly interferes with the stages of cell cycle and target caspase specific cell death (Antonsson & Persson, 2009; Shirmanova et al., 2017; Tawfik et al., 2017). We reveal SatIII functions antagonist to the growth inhibitory drugs. A null to diminished SatIII expression in the still-alive cells suggests an existing underlined repair mechanism that triggers interplay between SatIII transcripts and the administrated drug. This understanding gains a line of support with report stating new way of chemo-therapy leaves such still-alive cells (cells still undergoing DNA repair, or at pre-cancerous stage) with enough duration to acquire mutations that limit their toxic effect within one to few organs. This in turn helps a second line of treatment to target only the affected portion to avoid killing the normal dividing cells (Liu et al., 2015). Heat stress combined with anti-cancer drug (Tiligada, 2006) altogether increased the level of SatIII transcription. One possible explanation could be that the heat shock releases abundant heat shock proteins in tumor cells to confer protection against several stressful conditions (*reviewed in* Liu et al., 2015). The tumor derived-HSPs themselves boost up tumor immunity and hence support uncontrolled growth, the reason anti-cancer therapies do not prefer to raise the level of HSPs while targeting cell growth arrest (Tiligada, 2006; Liu et al., 2015). Since the apoptosis signals are bypassed in many cancer cells, the lncRNAs are often overexpressed or downregulated in order to sensitize tumor cells to various death stimuli (*reviewed in* Pecero et al., 2019). The drugs while inducing damage to the tumor cells sometimes may incorporate DNA lesion to the surrounding normal cells. This immediately activates DNA repair pathways in both the normal and the targeted tumor cells. The damage if rigorously repaired, often makes the cells resistant to the given chemo therapy (Kiwerska & Szyfter, 2019). Interestingly, 5-FU and cisplatin are among those cytotoxic drugs, against which chemoresistance can be efficiently developed (*reviewed in* Kiwerska & Szyfter, 2019). Since, our assays are limited to cell survival analysis, we show forced expression of SatIII prevents cell vulnerability towards anti-cancer drugs and support malignant growth. Several modifiers are involved to make cells susceptible to 5-FU. Among them, downregulation of lncRNAs is a good example of moderating drug-tolerance. Our result is in line and suggests that downregulation of oncogenic SatIII enhances chemosensitivity of the cancer cells. With overall observations, we therefore suggest SatIII RNA protects cancer cells form chemo-induced death. This, uncovers a possibility for the recent day’s strategy that targets components other than the DNA molecules (deactivating oncogenes or potentiating tumor suppressor genes), including the lncRNAs to prevent malignant transformation (Tiligada, 2006). Such targeted therapy is now referred as ‘non-oncogenic addiction’, and bears high prognostic value (Liu et al., 2015; Toma-Jonik et al., 2019). The drug-treated cells acquire activated damage repair mechanism, therefore, harbour functional p53 in order to promote cell cycle arrest (Upadhyay et al., 2016; Toma-Jonik et al., 2019). Interestingly, the tumor suppressing or tumor driving lncRNAs regulate (regulators) or are regulated (effectors) by p53 in order to control transcriptional activity of genes involved in cell cycle arrest and apoptosis (Sanchez Calle et al., 2018). It is tempting to demonstrate that in cancer cells SatIII colocalizes with p53 during the formation of nuclear stress bodies. Interestingly, SatIII knockdown leads to a higher expression of p53 protein. On the other hand, p53 can also regulate the level of SatIII transcription in order to induce cell death. The transcriptional status of both lncRNAs and p53 determines their function as pro- or anti-apoptotic elements, hence can be considered as crucial biomarkers for therapeutic targets (Chen et al., 2018). Our results here reveal, while SatIII restricts malignant cells from being susceptible to chemotherapeutic drugs, its knockdown restores functional p53 in the apoptotic cells. Although, we limit our study from looking into the molecular interactions, one line of evidence from recent reports suggests, reactivation of functional Tp53 in cancer cells sensitizes them towards chemotherapy, thus, increases treatment benefits (Bossi & Sacchi, 2007). In future, we are interested to identify the mechanism of action through which SatIII regulates different oncogenic events. This study establishes SatIII is an oncogenic lncRNA as it confers a pro-cell survival function and protects cancer cells from stress induced-damage. Therefore, a novel insight into the specific role of SatIII transcripts in cancer cell survival is being deciphered by this study.

## List of Abbreviations

LncRNAs: long non-coding RNAs
SatIII: satellite III
HSF1: heat shock factor 1
RNP: ribonucleoprotein
nSBs: nuclear stress bodies
SF: splicing factor
SRp: serine and arginine-rich protein
CREBBP: CREB (cAMP-response element binding protein) binding protein
hnRNPs: heterogeneous nuclear RNPs
HSR: heat shock response
HSPs: heat shock proteins
HOTAIR: HOX transcript antisense RNA
HOX: homeobox gene
MALAT1: metastasis associated lung adenocarcinoma transcript 1
PANDA: P21-associated ncRNA DNA damage-activated
PVT1: plasmacytoma variant translocation 1
GAS5: growth arrest specific 5
MEG3: maternally expressed 3
TonEBP: tonicity enhancer-binding protein
CDKs: cyclin-dependent kinases
5-FU: 5-fluorouracil
cisplatin: (cis-Diamminedichloroplatinum (II): CDDP)
DMEM: Dulbecco’s modified Eagle’s medium
RPMI: Roswell Park Memorial Institute medium
FBS: fetal bovine serum
FISH: fluorescence *in situ* hybridization
shRNA, MTT: 3-(4,5-dimethylthiazol-2-yl)-2,5-diphenyltetrazolium bromide
DMSO: dimethyl sulfoxide
NEAT1: nuclear paraspeckle assembly transcript 1
SWI/SNF: SWItch/Sucrose Non-Fermentable
BRD4: bromodomain-containing protein 4
UCA1: urothelial carcinoma-associated 1
PI3K/Akt: phosphatidylinositol-3-kinase/protein kinase B
NF-κB: nuclear factor-κB.

## ACKNOWLEDGEMENTS

The authors thank Prof. S. Ganesh for his scholarly inputs, valuable time, suggestions and advices to improve the study. We are also immensely thankful to Prof. S. Ganesh for providing his laboratory facilities at the Dept. of Biological Sciences and Bioengineering, IIT Kanpur. We thank Dr. Rashmi Parihar for her valuable involvement throughout the study. We also thank Prof. S. C. Lakhotia for his comments and suggestion throughout the study. Authors would like to thank the anonymous referees for valuable comments which helped to improve the manuscript. Authors thank Vellore Institute of Technology, Vellore for providing SSG with “VIT Seed Grant” for carrying out this research work.

## DATA AVAILABILITY STATEMENT

The data sets used and/or analyzed during the current study are available from the corresponding author on reasonable request.

## FIGURE LEGENDS

***Supplementary-SI***

**FIGURE S1** Treatment standardization and quantification of cell survival data. The treatment dosage and duration was determined by MTT analysis and the change in fold survival was noted. **a** HeLa was treated with 1μM, 2μM staurosporine, 0.2mM, 0.4mM 5-FU and 2.4μM, 4.8μM cisplatin for 12, 24, 48h durations and was subjected to MTT without heat shock. Increased cell death was observed with increase in the treatment dosage and durations; 24h of treatment was finalized with the minimum concentration of each drug and were either heat shocked, or kept as is at 37°C prior to MTT assay. The optimum percentage of cell survival was required to process the cells for *in situ* hybridization (**b**). **C** Similarly, HCT-15 was incubated with increasing concentrations of staurosporine and 5-FU and was further subjected to MTT analysis at three different time points. As shown in (**d**), the fold change in cell survival without or with heat shock was calculated against the non-treated control cells. DMSO was used as vehicle. NHS=no heat shock, HS=heat shock. Unpaired Student’s *t*-test, P ** ≤ 0.005, *** ≤ 0.0005. Scale bar: 10μM

***Supplementary-SII***

**FIGURE S2** RT-PCR result for SatIII overexpression and knockdown approach. Gel image represents the relative expression levels of SatIII transcripts in HeLa and HCT-15 cells as measured by a semi-quantitative RT-PCR. pcDNA is a mammalian expression vector harbouring SatIII repeat, thus used as control. Overexpression of SatIII resulted intense bands; reduced levels of SatIII transcript was noted in SatIII knocked down cells. SatIII sense=overexpressed transcripts. SatIII antisense=knocked down transcripts

