## Supplementary figures and images for "Human Satellite III long non-coding RNA imparts survival benefits to cancer cells"

### Supplementary Fig 1

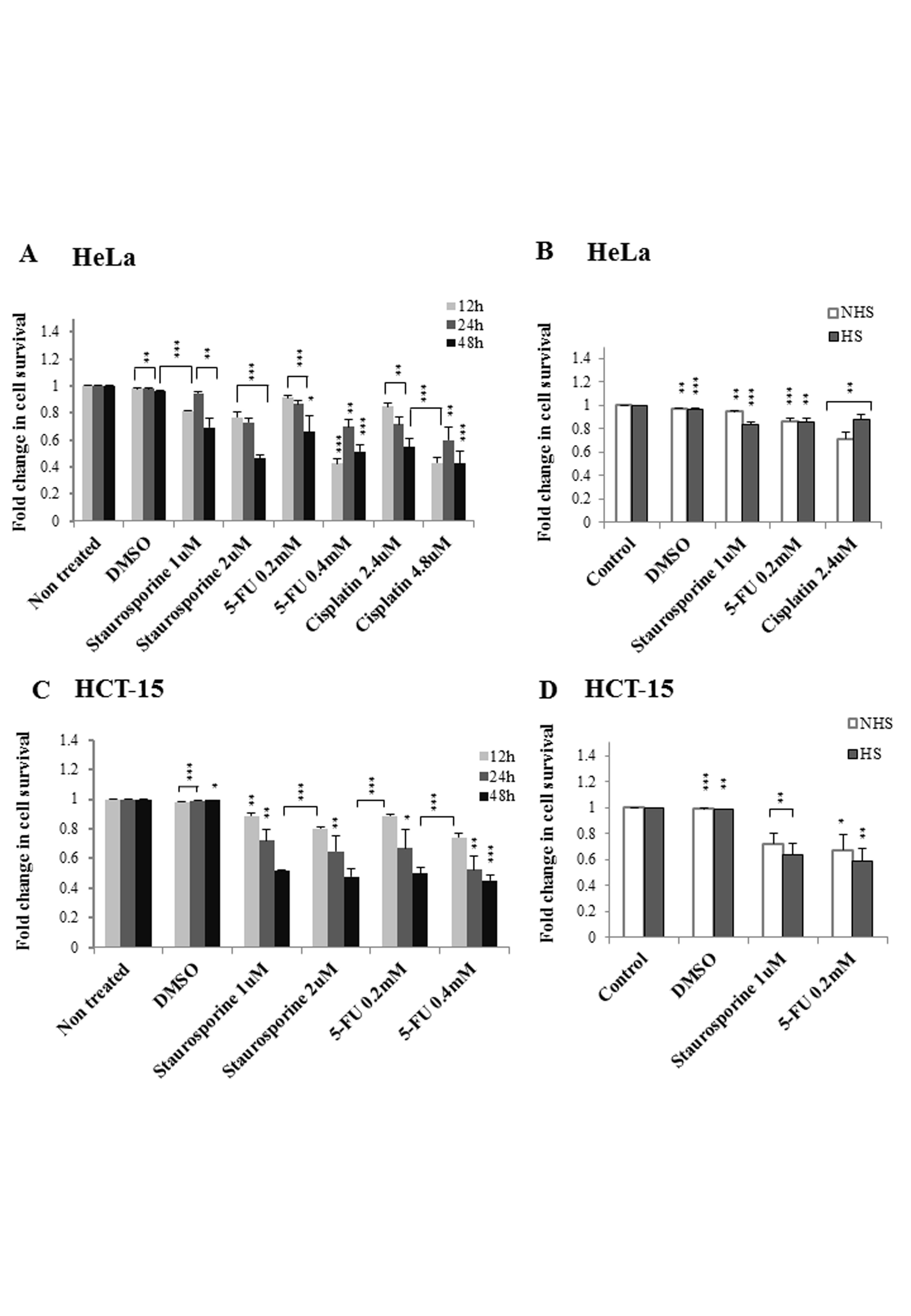

### Supplementary Fig 2

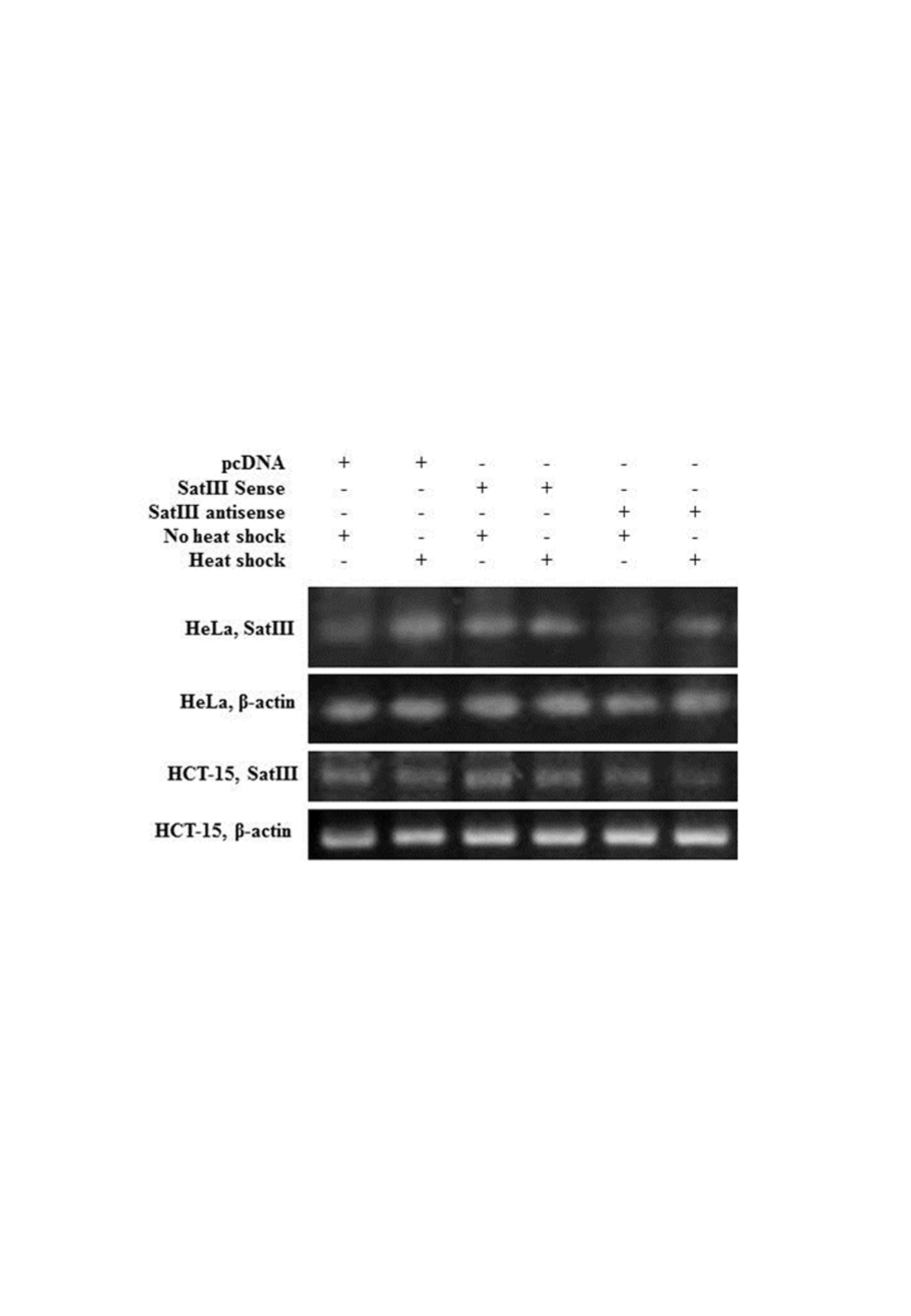
